# Identifying fundamental properties of neuromuscular disorder genes through genomic feature analysis

**DOI:** 10.64898/2026.04.21.719902

**Authors:** Alexandria Martin, M. Alejandra Llanes-Cuesta, Jessica N. Hartley, Patrick Frosk, Britt I. Drögemöller, Galen E.B. Wright

**Affiliations:** Department of Pharmacology and Therapeutics, Rady Faculty of Health Sciences, University of Manitoba, Winnipeg, Canada; Department of Biochemistry and Medical Genetics, Rady Faculty of Health Sciences, University of Manitoba, Winnipeg, Canada; PrairieNeuro Research Centre, Kleysen Institute for Advanced Medicine, Health Sciences Centre and Rady Faculty of Health Sciences, University of Manitoba, Winnipeg, Canada; Children’s Hospital Research Institute of Manitoba, Winnipeg, Canada; Department of Pediatrics and Child Health, Rady Faculty of Health Sciences, University of Manitoba, Winnipeg, Canada

**Author notes:** Correspondence to: Galen E.B. Wright, Department of Pharmacology and Therapeutics, University of Manitoba, Winnipeg, MB R3E 0T6, Canada.

## Abstract

Neuromuscular disorders (NMDs) encompass conditions that affect the peripheral nervous system. They are often caused by genetic alterations that impair skeletal muscle function. Molecular diagnosis remains challenging because some causal genes have not yet been identified. We therefore used feature-selection approaches to characterize genomic properties that distinguish NMD genes from other protein-coding genes.

Curated genes implicated in NMDs (*n* = 639; GeneTable of NMDs) were merged with 134 genomic features for autosomal protein-coding genes spanning gene complexity, genetic variation, expression patterns, and other gene traits. Machine-learning-based feature selection using Boruta was combined with statistical analyses to compare NMD genes with other protein-coding genes. We also evaluated NMD genes against high-confidence non-NMD disease genes, examined variation across NMD subgroups, and assessed genes added to the NMD catalogue between 2020 and 2024.

Compared with other protein-coding genes, NMD genes showed higher expression in disease-relevant tissues, including skeletal muscle and heart; greater transcript complexity and population genetic constraint, and stronger conservation across several evolutionary measures. Of 33 features that distinguished NMD genes from both high-confidence non-NMD disease genes and other autosomal genes, 28 were expression measures. Feature profiles varied across NMD subgroups, while genes added between 2020 and 2024 showed profiles similar to established NMD genes across all Boruta-confirmed features.

This study identified genomic features that distinguish NMD genes from high-confidence disease genes and broader autosomal protein-coding genes. These findings provide insight into the gene architecture, expression, constraint, and evolutionary conservation of NMD genes and may inform candidate-gene prioritization in genetically unresolved NMDs.

## Introduction

Neuromuscular disorders (NMDs) are a diverse group of conditions that affect the peripheral nervous system, including motor neurons, neuromuscular junctions, and skeletal muscle (1). Globally, an estimated 15 million people are affected by NMDs, corresponding to a prevalence of approximately one in 1,000 individuals, though this rate varies by region (2–4). The majority of NMDs have a genetic etiology, and advances in sequencing technologies have led to the identification of over 600 genes that are known to cause NMDs (5,6).

Despite this progress, the diagnostic yield for these conditions remains below 60% (7,8), and the continued expansion of the NMD disease-gene catalogue indicates that additional causal genes remain to be identified. Many individuals remain undiagnosed due to limitations of available gene panels and exome sequencing, the complexity of variant interpretation, and the presence of causal variants in genes and genomic regions not yet linked to NMDs (7,9). As a result, affected individuals often face a prolonged “diagnostic odyssey,” which can delay access to targeted interventions, clinical support resources, and genetic counselling (10,11). Diagnostic strategies that span larger genomic regions, such as whole-genome sequencing, are increasingly being applied for unresolved cases (12). These approaches employ computational filtering techniques to prioritize candidate variants, but are limited by the large number of detected variants and the difficulty of distinguishing pathogenic variants from benign background variation (13). Since the human genome contains nearly 20,000 protein-coding genes, systematic approaches to identify previously unknown causal NMD genes are crucial for informing genetic diagnoses and for understanding disease mechanisms and neurobiology (14).

An approach for improving disease gene discovery is to analyze annotated genomic features to identify shared biological properties that distinguish disease-associated genes from the broader genome (15–18). In recent years, multi-omic annotations have become increasingly available due to technological and methodological advances, enabling comprehensive exploration of gene properties (9,16). Thus, using these features can aid in prioritizing novel candidate disease genes and enhance the interpretation of gene-phenotype relationships (18). This strategy has become more feasible with the widespread availability of public resources, such as the GeneTable of NMDs, which catalogues the growing number of genes robustly linked to these disorders (5).

Similar computational approaches have been applied to more restricted disease groups, highlighting the need to extend these analyses to broader disease contexts. For example, an investigation used multi-omic annotations to characterize genes implicated in hereditary ataxia, a subtype of NMDs (9). Using computational approaches, the study revealed shared patterns across childhood- and adult-onset ataxia genes, including underappreciated contributors such as short tandem repeat (STR) density. However, these methods have not been applied across the larger set of NMD genes, creating an important knowledge gap.

Machine-learning-based feature selection provides one means of integrating these functional genomic annotations. Boruta is a random forest-based feature-selection algorithm. A random forest combines many decision trees, each built from a different sample of the data, to identify variables that consistently help distinguish groups. Boruta then compares each real variable with randomized *“shadow”* versions to determine whether it is more informative than expected by chance (19). Boruta has previously been used in various human genetic studies, including identifying discriminative features associated with gain- and loss-of-function pathogenic variants, as well as prioritizing relevant genes from genetic variant data in Alzheimer’s disease (20,21).

We therefore hypothesized that applying a genome-wide feature selection approach across all NMD subtypes could reveal shared biological features and provide insight into their genetic architecture. To investigate this, we compiled an extensive, curated list of genomic annotations from several public datasets and integrated them with a list of known NMD genes. We then applied Boruta feature selection along with complementary analytical approaches to identify the most important features for distinguishing NMD genes from other protein-coding genes (**Figure 1**). We further compared NMD genes with other high-confidence disease genes, examined variation across NMD subgroups, and assessed whether the identified feature patterns were also present among genes added to the NMD catalogue more recently. This analysis aimed to gain insight into the biological characteristics of NMD genes and establish a basis for testing the utility of these features in future candidate-gene prioritization studies.

**Figure 1.**
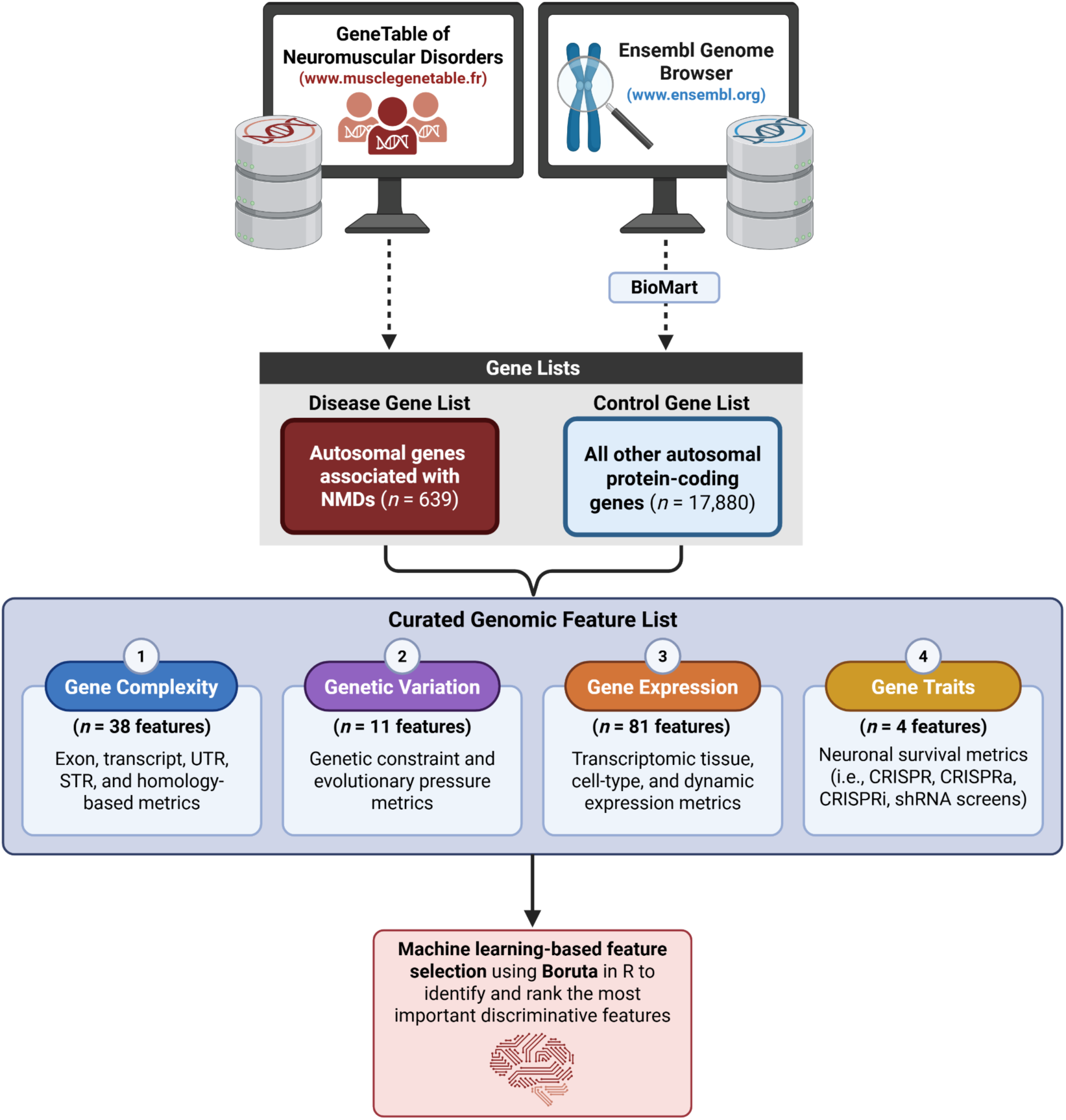
Overview of study design for identifying discriminative features of NMD genes. A list of genes associated with NMDs (*n* = 639) was obtained from the GeneTable of NMDs and merged with a curated genomic feature dataset for other human autosomal protein-coding genes (*n* = 17,880) from Ensembl. The two gene sets were evaluated across 134 genomic features within four categories: gene complexity (*n* = 38), genetic variation (*n* = 11), gene expression (*n* = 81), and gene traits (*n* = 4). Machine-learning-based feature selection using Boruta was applied to identify the most important features distinguishing NMD genes from other genes in the genome. Abbreviations: CRISPR, clustered regularly interspaced short palindromic repeats; CRISPRa, CRISPR activation; CRISPRi, CRISPR interference; NMDs, neuromuscular disorders; shRNA, short hairpin RNA; STR, short tandem repeat; UTR, untranslated region.

## Results

### Genomic features distinguish NMD genes from other autosomal protein-coding genes

We evaluated the discriminative potential of genomic features to distinguish NMD genes (*n* = 639) from other protein-coding genes (*n* = 17,880) using the Boruta machine-learning-based feature selection algorithm and complementary statistical techniques. Of the 134 features across four categories, Boruta classified 107 features (79.9%) as *“confirmed*,” indicating that they were important for distinguishing between the two gene lists (**Supplementary Figure S1**). This result indicates that differences between NMD genes and other autosomal protein-coding genes extend across multiple genomic feature classes.

To gain further insight into the features most informative for distinguishing NMD genes, we focused on the top 10 confirmed features ranked by median Boruta importance (**Figure 2A**). Boruta determines feature relevance by comparing observed importance scores with those of randomized shadow features, which were used in the analysis but are not shown in Figure 2. Skeletal muscle gene expression [quantified using the Genotype-Tissue Expression (GTEx) *t-*statistic] ranked as the most important feature, followed by gene expression in the heart left ventricle. Higher GTEx t-statistics indicate relatively greater expression in the indicated tissue. Other highly ranked features included heart *Tau* (tissue-specificity score), transcript count, evolutionary measures based on omega values for the rhesus macaque (*Macaca mulatta*) and marmoset (*Callithrix jacchus*), expression in the heart atrial appendage, mouse (*Mus musculus*) homolog percent identity, and measures of population genetic constraint based on gnomAD loss-of-function (LoF) and missense observed-to-expected (*o/e*) ratios. Boruta also *“rejected”* several features, including neuronal survival metrics, differentially developmentally regulated (DDG) values, expression in several less disease-relevant tissues (e.g., adrenal gland, thyroid, lung, etc.) and some region-specific STR measures.

**Figure 2.**
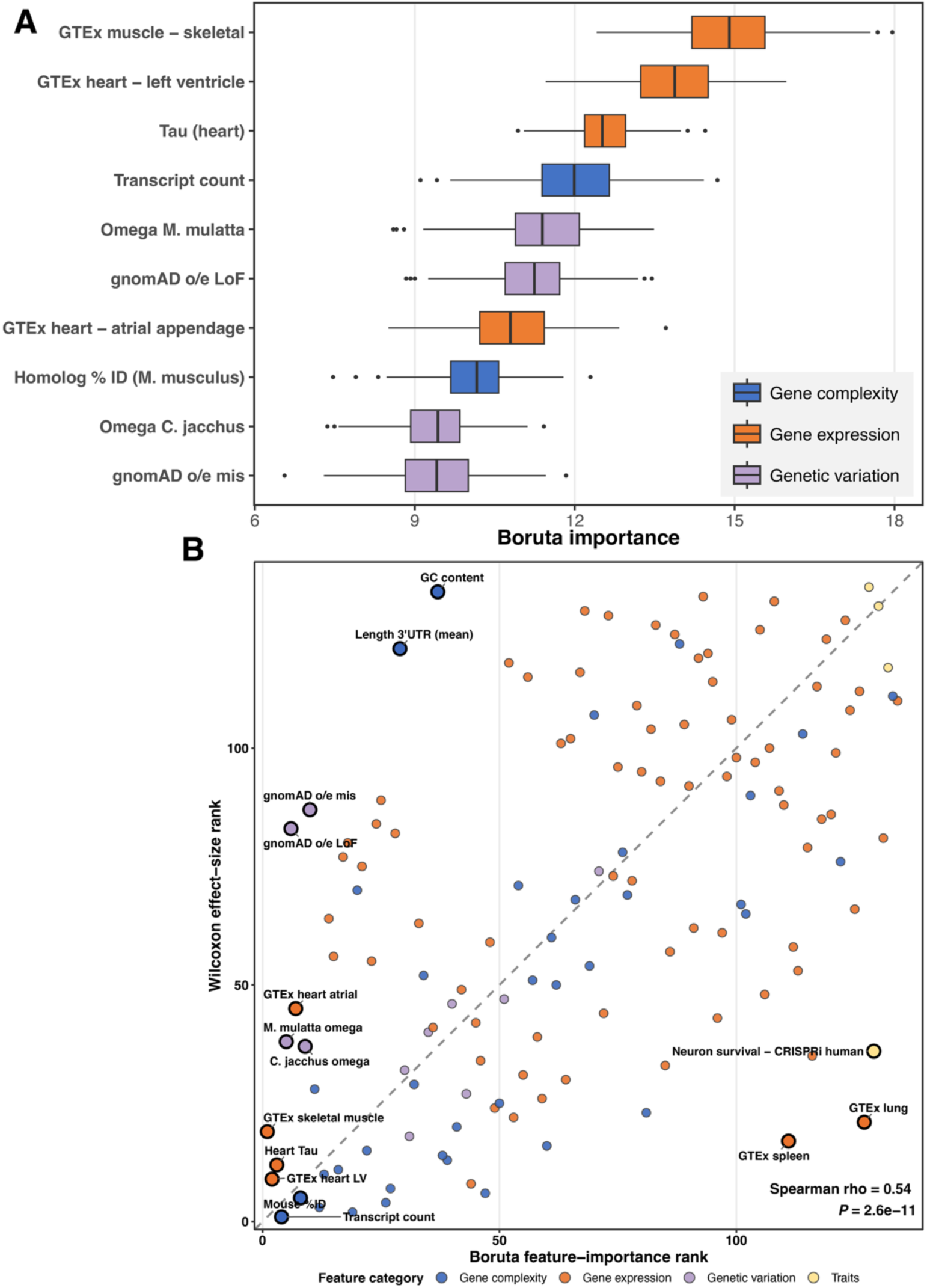
Boruta identifies genomic features that distinguish NMD genes, with concordance between Boruta importance and Wilcoxon effect-size rankings. **(A)** Boruta importance distributions across iterations for the 10 highest-ranked confirmed features distinguishing NMD genes (*n* = 639) from other autosomal protein-coding genes (*n* = 17,880). Features were ranked by median Boruta importance and are coloured by category: gene complexity (blue), gene expression (orange), and genetic variation (lavender). Randomized shadow features were included in the Boruta analysis but are not shown in the figure. Boxplots show the median and interquartile range; whiskers extend to 1.5 times the interquartile range, with more extreme values shown as points. **(B)** Comparison of Boruta feature-importance rank, based on median importance across iterations, with rank based on absolute Wilcoxon effect size for 134 genomic features. Colours indicate feature category. The 10 highest-ranked Boruta features and the five features with the largest differences between the two rankings are labelled. Rankings were positively correlated (Spearman *ρ* = 0.54, *P* = 2.6 × 10^-11^). Abbreviations: GTEx, Genotype-Tissue Expression; LoF, loss-of-function; mis, missense; o/e, observed/expected; UTR, untranslated region.

We complemented the Boruta analysis by calculating Wilcoxon absolute effect sizes for each feature to quantify differences between NMD and other autosomal protein-coding genes, using Wilcoxon rank-sum tests with multiple-testing correction to assess significance (Bonferroni-adjusted *P-*value < 0.05; **Supplementary Figure S2**). Of the 134 features analyzed, 72 differed significantly between the gene lists. The features with the largest effect sizes included transcript count (absolute *r* = 0.097), several exon-count metrics (absolute *r* = 0.070-0.086), and maximum transcript length (absolute *r* = 0.076), all of which were higher in NMD genes. *Mus musculus* homology percent identity also had a relatively large effect size (*r* = 0.077) and was higher in NMD genes. Features with higher Boruta importance generally showed larger absolute effect sizes, indicating agreement between the machine-learning-based and traditional statistical analyses. We therefore compared the rankings of features by Boruta importance with their rankings by Wilcoxon absolute effect size (**Figure 2B**). Overall, we observed a modest positive correlation between Boruta importance rank and effect size rank (Spearman *ρ* = 0.54, *P* = 2.6 × 10^-11^). This indicates that the two analysis methods capture overlapping but non-identical aspects of the data. Top-ranked features, including GTEx skeletal muscle, GTEx heart left ventricle, and transcript count, ranked relatively highly in both analyses, while features such as GTEx lung, GTEx spleen, GC content and mean 3’UTR length were discordant between rankings.

### Comparison with other high-confidence disease genes identifies NMD-associated features

Because some genomic features associated with NMD genes may also be common to disease genes, we compared NMD genes with high-confidence disease genes from the Gene Curation Coalition (GenCC) (22). High-confidence GenCC genes were defined as those with *“definitive”* or *“strong”* gene-disease classifications, after excluding genes already included in the NMD set. This resulted in 639 NMD genes, 3,468 high-confidence GenCC non-NMD genes, and 14,412 other autosomal protein-coding genes (**Figure 3**). Across all features, 37 differed significantly between NMD and GenCC non-NMD genes after Benjamini-Hochberg correction (FDR < 0.05). To identify features that more specifically distinguished NMD genes from other disease genes, we additionally required that features differ significantly between NMD genes and other autosomal protein-coding genes and show a concordant direction of effect across both comparisons. Thirty-three features met these FDR-based criteria (**Supplementary Figure S3**).

**Figure 3.**
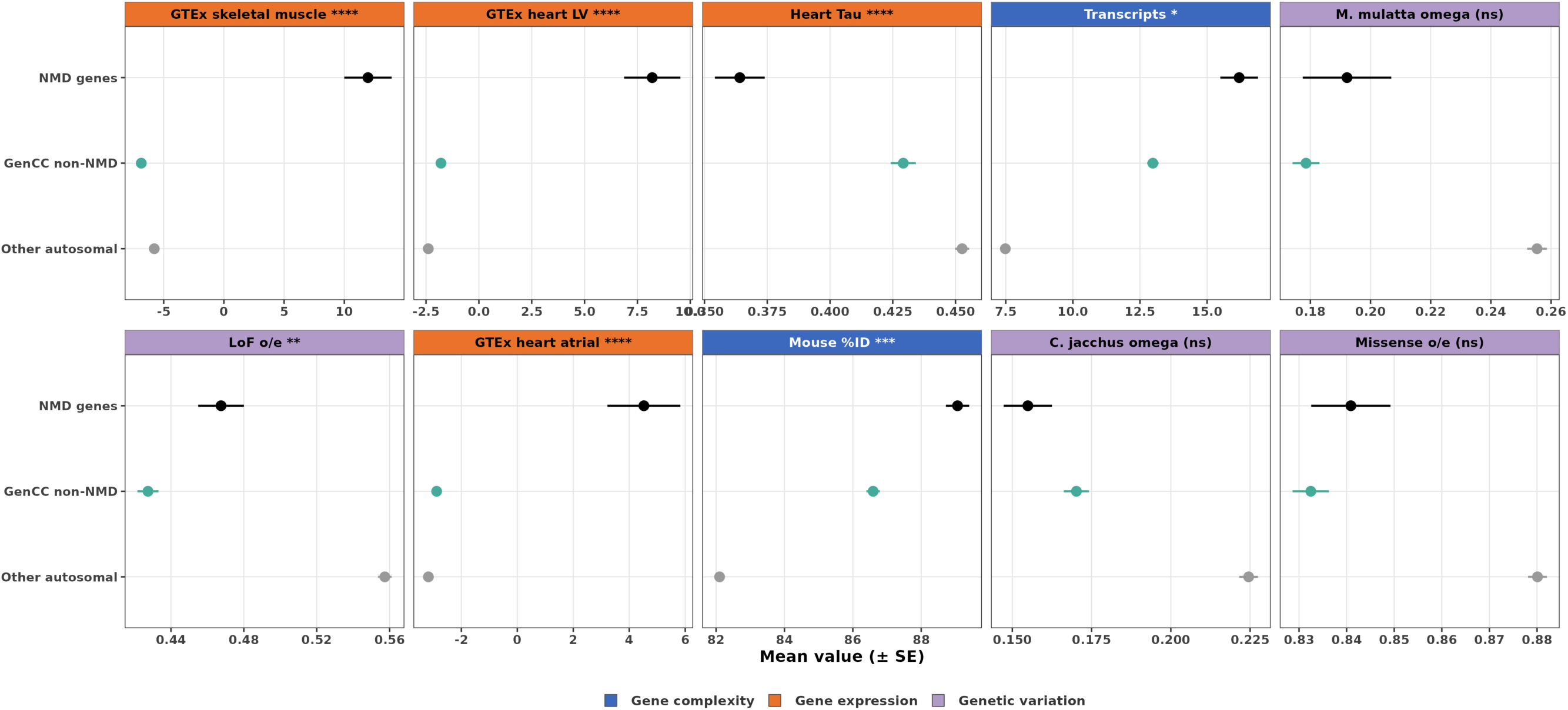
Comparison of top-ranked genomic features between NMD genes and other high-confidence disease genes. Mean values (± standard error, SE) are shown for the 10 highest-ranked Boruta features across NMD genes (*n* = 639), high-confidence GenCC disease genes not associated with NMDs (*n* = 3,468), and other autosomal protein-coding genes (*n* = 14,412). High-confidence GenCC genes were defined as those with *“definitive”* or *“strong”* gene-disease classifications after excluding NMD genes. Group sizes indicate the number of genes assigned to each group; plotted means and SEs use available values for each feature. Significance refers to Wilcoxon rank-sum comparisons between NMD and GenCC non-NMD genes, with Benjamini-Hochberg correction across features: *FDR < 0.05, **FDR < 0.01, ***FDR < 0.001, ****FDR < 0.0001; ns, not significant. Abbreviations: FDR; false discovery rate, GenCC; Gene Curation Coalition, GTEx, Genotype-Tissue Expression; LV, left ventricle; LoF, loss-of-function; mis, missense; o/e, observed/expected; SE, standard error.

Most were gene-expression measures (28/33), with four gene-complexity features and one gene-trait feature; no genetic-variation features met the NMD-distinguishing criteria.

We next examined the highest-ranked Boruta features in this comparison. GTEx skeletal muscle and heart left ventricle expression were higher in NMD genes compared to other high-confidence disease genes (effect size *r* = 0.130 and 0.133, respectively), as was heart atrial appendage expression (*r* = 0.089), whereas heart *Tau* was lower (*r* = -0.096) (**Figure 3**). Mouse homolog percent identity (*r* = 0.065) and transcript count (*r* = 0.042) were also significantly higher in NMD genes. In contrast, the primate *omega* measures and gnomAD missense *o/e* ratio did not differ significantly between NMD and GenCC non-NMD genes. The gnomAD LoF *o/e* ratio was higher in NMD genes than in GenCC non-NMD genes. These results indicate that tissue-expression measures comprised most of the features distinguishing NMD genes from other high-confidence disease genes, whereas several constraint and evolutionary features identified in the initial analysis are shared more broadly across disease-associated genes.

### Genomic features vary across NMD disease subgroups

Subgroup analyses showed that the top-ranked genomic features varied across NMD categories (**Figure 4**). The clearest differences involved tissue expression: skeletal muscle and cardiac expression were highest in myopathy and cardiomyopathy groups, whereas several neurological NMD groups (e.g., ataxias, paraplegias, neuropathies) showed lower relative values in these tissues. In contrast, transcript count was elevated across a broader range of categories, including both muscle and neurological disorders, suggesting that increased transcript complexity is not restricted to a single clinical subgroup. Constraint and primate evolutionary metrics showed smaller and less consistent differences across NMD subgroups and, consistent with the GenCC comparison, appeared to reflect broader disease-gene properties rather than features specific to individual NMD classes (**Figure 4**).

**Figure 4.**
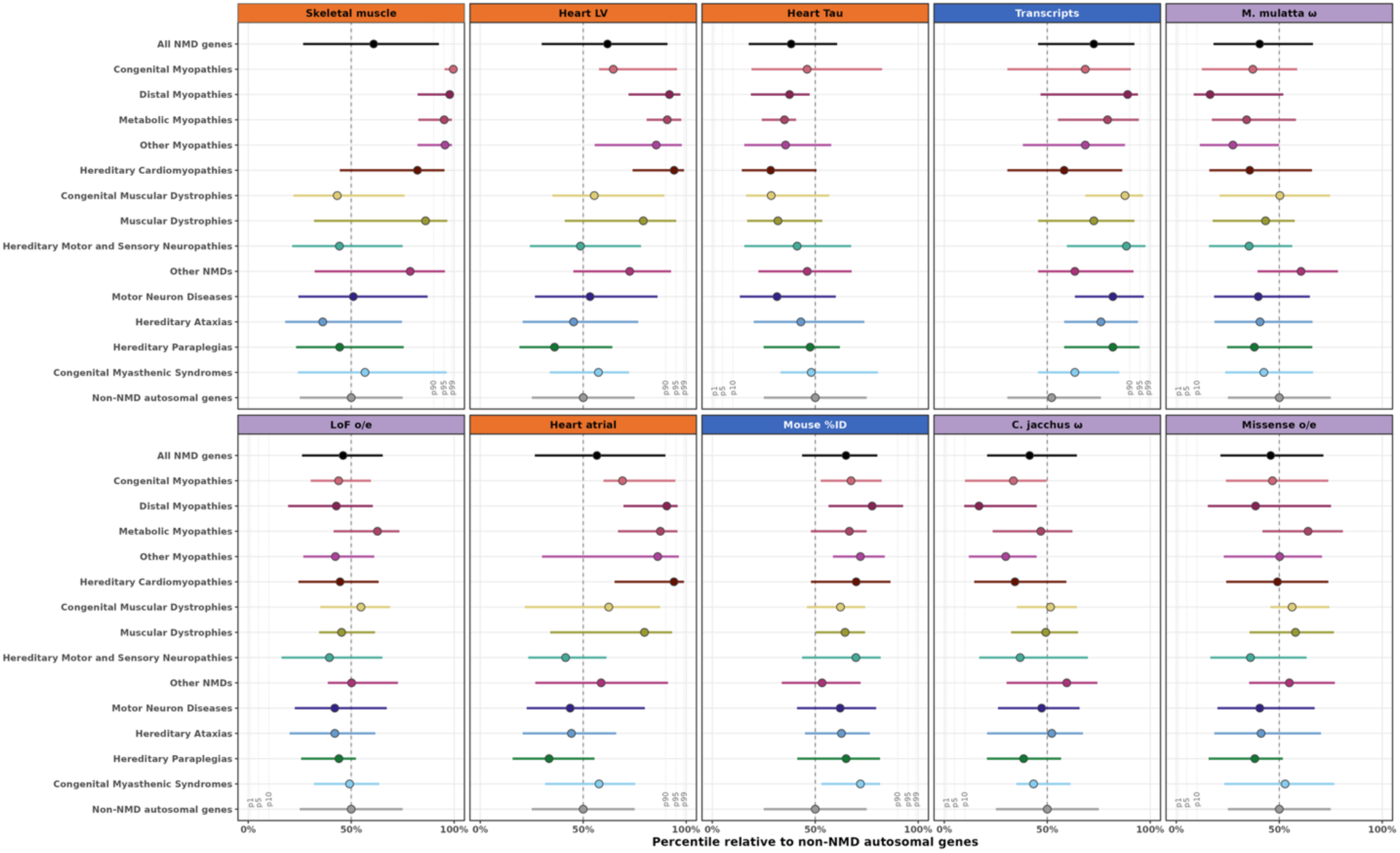
Top-ranked genomic features vary across NMD disease groups. Median feature percentiles and interquartile ranges are shown for the 10 highest-ranked Boruta features across 13 NMD disease groups containing at least 10 genes, all NMD genes, and non-NMD autosomal genes. Feature values were converted to percentiles relative to non-NMD autosomal genes. The dashed vertical line indicates the 50th percentile. Dotted lines indicate the 90th, 95th, and 99th percentiles for features with higher values in NMD genes and the 1st, 5th, and 10th percentiles for features with lower values in NMD genes. Direction was defined from the overall NMD versus non-NMD comparison. Genes associated with more than one NMD disease group were retained in each relevant group. Colours identify disease groups; all NMD genes are shown in black and non-NMD autosomal genes in grey. Abbreviations: GTEx, Genotype-Tissue Expression; LV, left ventricle; LoF, loss-of-function; mis, missense; o/e, observed/expected.

Selected expression, gene-complexity, and homology features are shown in greater detail in **Supplementary Figure S4**. Whole-blood expression was lower across several NMD groups, while transcript count, mean transcript length, and total STR count showed distinct subgroup distributions. Cross-species homology also varied by disease group. Mouse homolog percent identity was generally higher in NMD genes than in non-NMD genes, whereas zebrafish and *Drosophila* homology showed more heterogeneous patterns. Metabolic myopathy genes showed relatively high conservation in both zebrafish and *Drosophila*, while distal myopathy genes showed particularly low *Drosophila* homology. The proportion of genes with extreme NMD-like feature values also varied across disease groups, with subgroup-specific differences in the proportion of genes falling within the direction-specific 5% and 1% tails of the non-NMD distribution (**Supplementary Figure S5**).

### Genomic feature profiles are similar between established and more recently added NMD genes

To assess whether these genomic feature patterns were also present among more recently identified NMD genes, we compared genes established by July 2020 (*n* = 550) with genes added to the NMD gene list between 2020 and 2024 (*n* = 89). Median feature values, expressed as percentiles relative to other autosomal protein-coding genes, were positively correlated across all 107 Boruta-confirmed features (Spearman *ρ* = 0.77, *P* = 5.2 × 10^-22^) and across the top 10 features (*ρ* = 0.88, *P* = 8.1 × 10^-4^; **Figure 5A**). The top-ranked features showed similar profiles between established and more recently added NMD genes (**Figure 5B**; **Supplementary Figure S6**). These results indicate that the genomic feature patterns identified in established NMD genes are also observed in more recently added genes.

**Figure 5.**
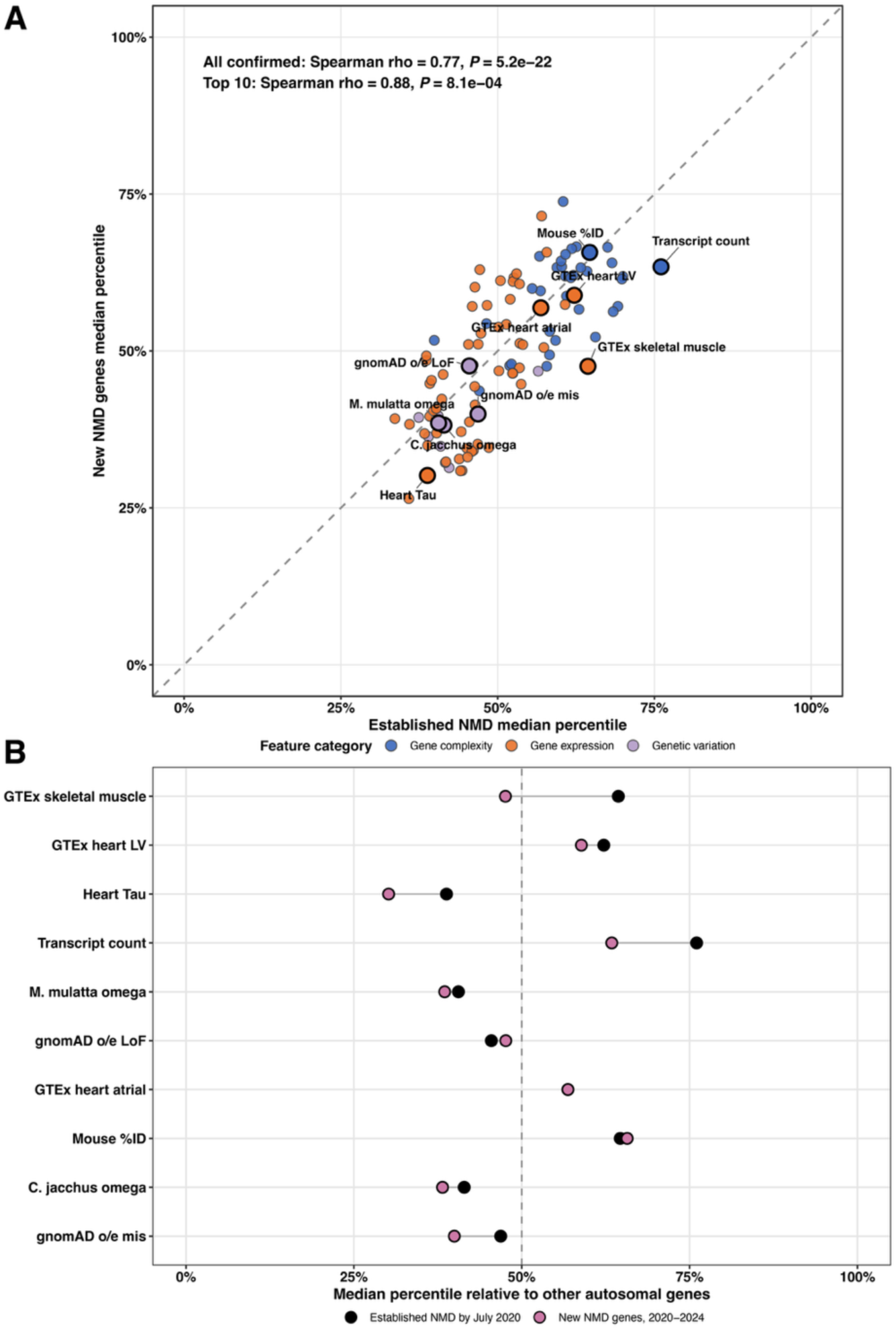
Genomic feature profiles are correlated between established and more recently added NMD genes. **(A)** Comparison of genomic feature profiles for NMD genes established by July 2020 (*n* = 550) and genes added to the NMD gene list between 2020 and 2024 (*n* = 89). For each feature, the median value within each gene set was expressed as a percentile relative to the distribution among other autosomal protein-coding genes. Points represent the 107 Boruta-confirmed features and are coloured by feature category; the 10 highest-ranked Boruta features are outlined and labelled. The dashed diagonal indicates equal percentiles between the two gene sets. Feature profiles were correlated across all confirmed features (Spearman ρ = 0.77, *P* = 5.2 × 10^-22^) and across the top 10 features (Spearman ρ = 0.88, *P* = 8.1 × 10^-4^). **(B)** Median feature values expressed as percentiles for the 10 highest-ranked Boruta features in established NMD genes (black) and genes added between 2020 and 2024 (rose). Values for the same feature are connected by grey lines; the dashed vertical line indicates the 50th percentile relative to other autosomal protein-coding genes. Abbreviations: GTEx, Genotype-Tissue Expression; LV, left ventricle; LoF, loss-of-function; mis, missense; o/e, observed/expected.

## Discussion

Our study identified genomic features that help distinguish NMD genes from other autosomal protein-coding genes and clarified which of these features are shared more broadly with disease-associated genes. Across 134 genomic features, NMD genes differed in disease-relevant tissue expression, gene and transcript complexity, population genetic constraint and cross-species conservation. Comparison with high-confidence GenCC disease genes separated properties shared more broadly across disease-associated genes from features that more specifically characterize NMD genes, which were dominated by tissue-expression measures. These patterns varied across NMD subgroups and were also evident among genes added to the NMD catalogue after 2020.

Tissue-expression measures were among the strongest discriminators of NMD genes, especially in tissues most directly affected by these disorders. Skeletal muscle and left ventricular heart expression ranked highest in the Boruta analysis and differed significantly between NMD and other autosomal protein-coding genes, whereas whole-blood expression was lower. The prominence of skeletal-muscle and cardiac expression is expected given the phenotypes represented in the NMD gene set, but these differences were also evident when NMD genes were compared with high-confidence GenCC non-NMD disease genes. Of the 33 features meeting the NMD-distinguishing criteria, 28 were expression measures, including higher expression in skeletal muscle and cardiac tissues. The subgroup analyses further showed that these patterns were not uniform across NMDs, with the strongest skeletal-muscle and cardiac profiles in myopathy and cardiomyopathy groups (23,24). These findings indicate that disease-relevant expression distinguishes NMD genes not only from the broader genome, but also from other established disease genes, while retaining clear differences across NMD subtypes.

Disease-relevant expression profiles can provide information for genetic variant interpretation that is not apparent from genome sequencing alone (25). When direct sampling of affected tissues is impractical, disease-relevant cellular models may provide complementary evidence for evaluating candidate genes and variants. In such cases, more accessible sources such as peripheral blood mononuclear cells or skin fibroblasts can be reprogrammed into patient-specific induced pluripotent stem cells (iPSCs), which can then be differentiated into relevant cell types or organoids to capture disease-specific gene expression profiles (26,27). This approach has already been employed in NMD research; for instance, studies have produced organoids from patient-derived iPSCs that serve as platforms for drug testing and recapitulating disease pathology in conditions such as muscular dystrophies and amyotrophic lateral sclerosis (28–31).

Gene-complexity measures were also prominent, with NMD genes tending to have more transcripts, longer transcripts and greater STR content than other autosomal genes. These properties are consistent with the involvement of large, extensively spliced genes such as *TTN* and *NEB,* and with the established contribution of alternative splicing to NMDs (32–37). NMD genes also had more annotated transcripts than high-confidence non-NMD disease genes, whereas transcript length and most exon-count measures showed less evidence of a direct difference between these groups. This suggests that isoform diversity, rather than gene size alone, may be a particularly relevant aspect of NMD gene architecture. Greater transcript diversity may increase the potential for tissue- or isoform-specific variant effects and supports isoform-aware annotation and transcript-resolved approaches such as long-read sequencing.

NMD subgroup analyses illustrated that these genomic feature patterns were not uniform. Transcript counts were elevated across a relatively broad range of muscle and neurological groups, whereas STR counts were highest in hereditary ataxias, hereditary motor and sensory neuropathies, hereditary paraplegias, and hereditary cardiomyopathies, with smaller shifts in most myopathy groups. Increased STR burden should therefore not be interpreted as a general property of every NMD category. Long-read DNA and RNA sequencing may be useful for resolving the transcript structures, isoforms, and repetitive sequences represented by these features (38).

Cross-species homology was also among the genomic features that distinguished NMD genes. Mouse homolog percent identity was among the top-ranked features and was higher in NMD genes than in other autosomal protein-coding genes. It also remained higher when NMD genes were compared with high-confidence GenCC disease genes. The retention of higher mouse homolog percent identity in the GenCC comparison indicates that this conservation signal is not explained entirely by properties shared across high-confidence disease genes. Homology patterns were less consistent in other commonly used NMD model organisms, including zebrafish and *Drosophila*, and varied across NMD subgroups. For example, metabolic myopathy genes showed relatively high conservation in both species, whereas distal myopathy genes showed particularly low *Drosophila* homology. These results suggest that sequence conservation should be considered when selecting model systems for individual candidate genes. In cases where conservation is limited, human cell-based systems, such as iPSCs, may provide a complementary platform for studying disease mechanisms (27).

The comparison of established and more recently added NMD genes also showed that these genomic patterns persisted as the NMD gene catalogue expanded. Feature profiles were strongly correlated between genes established by 2020 and those added between 2020 and 2024, both across all Boruta-confirmed features and among the top-ranked features. Although this retrospective analysis does not test prospective gene prediction, it indicates that the identified genomic characteristics are not restricted to the historically established NMD gene set.

While the analyses presented here offer insight into the biological characteristics of NMD genes, certain limitations should be considered. The comparison set of other autosomal protein-coding genes may include genes that are subsequently linked to NMDs or to incompletely characterized neuromuscular phenotypes. The GenCC analysis helps address this issue by comparing NMD genes with other high-confidence disease genes, although the underlying gene lists remain dependent on current disease annotations. Genes on sex chromosomes were excluded because several feature datasets were incomplete for these chromosomes, limiting the analysis of important X-linked NMD genes such as *DMD*, *MTM1*, and *EMD*. In addition, Boruta importance reflects the ability of a feature to distinguish gene sets rather than an independent or causal contribution to disease, and some features showed differences between Boruta and conventional effect-size rankings.

Future studies could extend the feature set as additional disease-relevant genomic and functional datasets become available. Genome-wide perturbation screens and long-read sequencing could add measures of gene function, transcript structure, isoform diversity, and repetitive sequence content that are not represented in the current analysis (38). Prospective studies will be needed to determine whether incorporating these features improves prioritization of previously unrecognized NMD genes.

In conclusion, NMD genes show a consistent genomic pattern characterized by disease-relevant tissue expression, gene complexity, and evolutionary conservation. Comparison with other high-confidence disease genes shows that several of these properties are shared more broadly across disease-associated genes, whereas tissue-expression features provide the clearest distinction of NMD genes. Subgroup analyses demonstrate substantial heterogeneity across NMD classes, but the overall feature profile is reproduced among genes added to the NMD catalogue more recently. These findings are limited to autosomal genes because sex-chromosome genes were excluded due to incomplete feature coverage. This consistency provides a basis for testing the value of these features in prospective candidate-gene prioritization studies. If validated in this setting, improved gene prioritization could contribute to higher molecular diagnosis rates in NMDs, with potential benefits for disease monitoring, treatment decisions, genetic counselling and access to support resources (11). The genomic features identified here may also provide additional context for the interpretation of genetic data in unresolved NMD cases.

## Materials and methods

### Database construction and NMD gene annotation

We built a comprehensive dataset of genomic features for human protein-coding genes. To ensure consistency across features, analyses were restricted to autosomal genes as several features were incomplete for sex chromosomes. Genes associated with NMDs were identified using the curated GeneTable of NMDs (https://musclegenetable.fr; accessed November 2024), resulting in 639 NMD genes and 17,880 other autosomal protein-coding genes. Genic information was obtained from Ensembl v.114 and the *biomaRt* package in R (v.2.58.2). Additional data-processing details are provided in Supplementary Methods.

### Construction of an extensive genomic feature list

To identify features that distinguish NMD genes, we assembled a set of 134 gene-level features for each gene from publicly accessible multi-omics resources, grouped into four major categories (**Figure 1**): (1) gene complexity (*n* = 38), (2) genetic variation (*n* = 11), (3) gene expression (*n* = 81), and (4) other gene traits (*n* = 4).

We obtained genomic features from publicly available resources, including measures of gene and transcript structure, repetitive sequence content, cross-species homology, population genetic constraint, evolutionary selection, tissue and developmental gene expression, and functional gene essentiality. Detailed definitions, processing procedures, and data sources for individual features are provided in **Supplementary Methods** and **Supplementary Table 1**.

All data sources were merged into a unified genomic feature list. Missing values were imputed using the *MICE* R package (v.3.17.0), using predictive mean matching to generate five imputed datasets (*m* = 5). The imputed feature matrix was used for Boruta feature selection, whereas statistical comparisons, subgroup analyses, GenCC comparisons, and temporal analyses used the original feature values, with missing values excluded on a feature-specific basis.

### Statistical analysis

For each genomic feature, we compared NMD genes with other autosomal protein-coding genes using Wilcoxon rank-sum tests with Bonferroni correction for multiple testing. Wilcoxon effect sizes (*r*) and corresponding 95% confidence intervals were calculated to quantify differences between groups. Effect size direction was assigned according to the direction of the mean differences between groups. Agreement between feature rankings based on Boruta importance and absolute Wilcoxon effect size was assessed using Spearman rank correlation and Kendall rank correlation. All statistical analyses were performed in R (v.4.3.3).

### Boruta feature selection

To identify genomic features that most strongly distinguish NMD genes from other autosomal protein-coding genes, we applied the Boruta algorithm (19) to the imputed feature matrix. Feature selection was performed using the R *Boruta* package (v.8.0.0) with a maximum of 250 iterations and a fixed random seed for reproducibility. Tentative feature classifications were then resolved using the *TentativeRoughFix()* function. Feature importance was summarized across Boruta iterations, and confirmed features were ranked by median importance for selection of the 10 highest-ranked features.

### Comparison with high-confidence disease genes

To determine whether genomic features associated with NMD genes were also characteristic of disease genes more generally, we compared NMD genes with high-confidence disease-associated genes from the GenCC (22). GenCC entries classified as *“definitive”* or *“strong”* were retained, and genes already present in the NMD set were excluded from the GenCC comparator group. Genes were assigned to three mutually exclusive groups: NMD genes (*n* = 639), high-confidence GenCC non-NMD genes (*n* = 3,468), and remaining autosomal protein-coding genes (*n* = 14,412).

For each genomic feature, pairwise Wilcoxon rank-sum tests were performed for NMD versus GenCC non-NMD genes, NMD versus other autosomal genes, and GenCC non-NMD versus other autosomal genes. Benjamini-Hochberg correction was applied separately within each comparison, with FDR < 0.05 considered significant. Wilcoxon effect sizes were assigned a direction based on the mean difference between groups. Features were classified as NMD-distinguishing when they differed significantly between NMD and GenCC non-NMD genes and between NMD and other autosomal genes, with concordant directions of effect in both comparisons.

### NMD disease subgroup analyses

To assess variation across NMD disease groups, genes were stratified according to the disease categories defined in the GeneTable of NMDs. Groups containing at least 10 genes (**Supplementary Table 2**) were retained (*n* = 13), and genes associated with more than one disease group were retained in each relevant group. For the top-ranked Boruta features and selected additional features, individual gene values were converted to percentiles relative to the corresponding distribution among non-NMD autosomal genes. Disease-group distributions were summarized using the median percentile and interquartile range.

To examine the contribution of genes with extreme feature values, thresholds were defined from the non-NMD distribution. Feature direction was determined from the overall NMD-versus-non-NMD comparison. For features with higher values in NMD genes, the NMD-like 5% and 1% tails corresponded to values above the 95^th^ and 99^th^ percentiles of non-NMD genes; for features with lower values in NMD genes, the corresponding tails comprised values below the 5^th^ and 1^st^ percentiles. The number and proportion of genes within these direction-specific tails were calculated for each disease group.

### Comparison of established and more recently added NMD genes

To assess whether the identified genomic feature patterns were also present among more recently identified NMD genes, we compared the GeneTable of NMDs from July 2020 with the November 2024 version used for the primary analysis. After restricting to autosomal genes represented in the feature dataset and harmonizing updated gene symbols, genes present in the 2020 list were classified as established NMD genes (*n* = 550), while genes present in the 2024 list but absent from the 2020 list were classified as more recently added NMD genes (*n* = 89).

For each of the 107 Boruta-confirmed features, the median feature value within each NMD gene set was converted to a percentile relative to the distribution among other autosomal protein-coding genes. Similarity between established and more recently added NMD genes was assessed using Spearman correlation across all confirmed features and separately across the 10 highest-ranked Boruta features. Mean values and standard errors for the top-ranked features were also calculated for visualization.

## Supporting information

Supplementary Information Other

## Acknowledgements

We gratefully acknowledge the data providers whose contributions, through open-access repositories or direct data sharing, made this genomic feature analysis possible. Limited sections of the manuscript and analysis code were edited for language clarity and code quality with assistance from ChatGPT (OpenAI, San Francisco, CA), without affecting the scientific content.

## Author contributions

A.M.: Data curation, Formal analysis, Investigation, Methodology, Visualization, Writing: original draft, Writing: review and editing. M.A.L.-C.: Data curation, Formal analysis, Project administration, Writing: review and editing. J.N.H.: Conceptualization, Investigation, Writing: review and editing. P.F.: Conceptualization, Supervision, Writing: review and editing. B.I.D.: Conceptualization, Funding acquisition, Methodology, Supervision, Writing: review and editing. G.E.B.W.: Conceptualization, Data curation, Formal analysis, Funding acquisition, Investigation, Methodology, Resources, Supervision, Visualization, Writing: original draft, Writing: review and editing.

## Data availability

The data analyzed in this study were obtained from publicly available resources described in the Materials and Methods and Supplementary Table 1. Analysis code will be deposited in a public repository and made available prior to publication.

## References

1. Dowling, J.J., D Gonorazky, H., Cohn, R.D. and Campbell, C. (2018) Treating pediatric neuromuscular disorders: The future is now. Am J Med Genet A, 176, 804–841.

2. Deenen, J.C.W., Horlings, C.G.C., Verschuuren, J.J.G.M., Verbeek, A.L.M. and van Engelen, B.G.M. (2015) The Epidemiology of Neuromuscular Disorders: A Comprehensive Overview of the Literature. J Neuromuscul Dis, 2, 73–85.

3. Wilson, L.A., Macken, W.L., Perry, L.D., Record, C.J., Schon, K.R., Frezatti, R.S.S., Raga, S., Naidu, K., Köken, Ö.Y., Polat, I., et al. (2023) Neuromuscular disease genetics in under-represented populations: increasing data diversity. Brain, 146, 5098–5109.

4. Zatz, M., Passos-Bueno, M.R. and Vainzof, M. (2016) Neuromuscular disorders: genes, genetic counseling and therapeutic trials. Genet Mol Biol, 39, 339–348.

5. Benarroch, L., Bonne, G., Rivier, F., Procaccio, V. and Hamroun, D. (2025) The 2025 version of the gene table of neuromuscular disorders (nuclear genome). Neuromuscul Disord, 46, 105261.

6. Laing, N.G. (2012) Genetics of neuromuscular disorders. Critical Reviews in Clinical Laboratory Sciences, 49, 33–48.

7. Koczwara, K.E., Lake, N.J., DeSimone, A.M. and Lek, M. (2022) Neuromuscular disorders: finding the missing genetic diagnoses. Trends Genet, 38, 956–971.

8. Yubero, D., Natera-de Benito, D., Pijuan, J., Armstrong, J., Martorell, L., Fernàndez, G., Maynou, J., Jou, C., Roldan, M., Ortez, C., et al. (2021) The Increasing Impact of Translational Research in the Molecular Diagnostics of Neuromuscular Diseases. IJMS, 22, 4274.

9. Chen, Z., Tucci, A., Cipriani, V., Gustavsson, E.K., Ibañez, K., Reynolds, R.H., Zhang, D., Vestito, L., García, A.C., Sethi, S., et al. (2023) Functional genomics provide key insights to improve the diagnostic yield of hereditary ataxia. Brain, 146, 2869–2884.

10. Bauskis, A., Strange, C., Molster, C. and Fisher, C. (2022) The diagnostic odyssey: insights from parents of children living with an undiagnosed condition. Orphanet J Rare Dis, 17, 233.

11. Dratch, L., Azage, M., Baldwin, A., Johnson, K., Paul, R.A., Bardakjian, T.M., Michon, S.-C., Amado, D.A., Baer, M., Deik, A.F., et al. (2024) Genetic testing in adults with neurologic disorders: indications, approach, and clinical impacts. J Neurol, 271, 733–747.

12. Mitsuhashi, S. and Matsumoto, N. (2020) Long-read sequencing for rare human genetic diseases. J Hum Genet, 65, 11–19.

13. Najafi, A., Caspar, S.M., Meienberg, J., Rohrbach, M., Steinmann, B. and Matyas, G. (2020) Variant filtering, digenic variants, and other challenges in clinical sequencing: a lesson from fibrillinopathies. Clinical Genetics, 97, 235–245.

14. Nurk, S., Koren, S., Rhie, A., Rautiainen, M., Bzikadze, A.V., Mikheenko, A., Vollger, M.R., Altemose, N., Uralsky, L., Gershman, A., et al. (2022) The complete sequence of a human genome. Science, 376, 44–53.

15. Capanu, M. and Ionita-Laza, I. (2015) Integrative analysis of functional genomic annotations and sequencing data to identify rare causal variants via hierarchical modeling. Front Genet, 6, 17.

16. Ionita-Laza, I., McCallum, K., Xu, B. and Buxbaum, J.D. (2016) A spectral approach integrating functional genomic annotations for coding and noncoding variants. Nat Genet, 48, 214–220.

17. Li, X., Li, Z., Zhou, H., Gaynor, S.M., Liu, Y., Chen, H., Sun, R., Dey, R., Arnett, D.K., Aslibekyan, S., et al. (2020) Dynamic incorporation of multiple in silico functional annotations empowers rare variant association analysis of large whole-genome sequencing studies at scale. Nat Genet, 52, 969–983.

18. Reynolds, R.H., Hardy, J., Ryten, M. and Gagliano Taliun, S.A. (2019) Informing disease modelling with brain-relevant functional genomic annotations. Brain, 142, 3694–3712.

19. Kursa, M.B. and Rudnicki, W.R. (2010) Feature Selection with the **Boruta** Package. J. Stat. Soft., 36.

20. Ahmed, H., Soliman, H. and Elmogy, M. (2022) Early detection of Alzheimer’s disease using single nucleotide polymorphisms analysis based on gradient boosting tree. Comput Biol Med, 146, 105622.

21. Sevim Bayrak, C., Stein, D., Jain, A., Chaudhary, K., Nadkarni, G.N., Van Vleck, T.T., Puel, A., Boisson-Dupuis, S., Okada, S., Stenson, P.D., et al. (2021) Identification of discriminative gene-level and protein-level features associated with pathogenic gain-of-function and loss-of-function variants. The American Journal of Human Genetics, 108, 2301–2318.

22. DiStefano, M.T., Goehringer, S., Babb, L., Alkuraya, F.S., Amberger, J., Amin, M., Austin-Tse, C., Balzotti, M., Berg, J.S., Birney, E., et al. (2022) The Gene Curation Coalition: A global effort to harmonize gene-disease evidence resources. Genet Med, 24, 1732–1742.

23. Goodwin, J.F. (1992) Cardiomyopathies and specific heart muscle diseases. Definitions, terminology, classifications and new and old approaches. Postgrad Med J, 68 S1, 3–6.

24. Nagy, H. and Veerapaneni, K.D. (2026) Myopathy. StatPearls: GeneReviews, Treasure Island (FL).

25. Cummings, B.B., Marshall, J.L., Tukiainen, T., Lek, M., Donkervoort, S., Foley, A.R., Bolduc, V., Waddell, L.B., Sandaradura, S.A., O’Grady, G.L., et al. (2017) Improving genetic diagnosis in Mendelian disease with transcriptome sequencing. Sci Transl Med, 9, eaal5209.

26. El Hokayem, J., Cukier, H.N. and Dykxhoorn, D.M. (2016) Blood Derived Induced Pluripotent Stem Cells (iPSCs): Benefits, Challenges and the Road Ahead. J Alzheimers Dis Parkinsonism, 6, 275.

27. Speciale, A.A., Ellerington, R., Goedert, T. and Rinaldi, C. (2020) Modelling Neuromuscular Diseases in the Age of Precision Medicine. JPM, 10, 178.

28. Auletta, B., Chiolerio, P., Cecconi, G., Rossi, L., Sartore, L., Cecchinato, F., Barbato, G., Lauroja, A., Maghin, E., Easler, M., et al. (2025) Tissue-engineered neuromuscular organoids. Commun Biol, 8, 1074.

29. Cheesbrough, A., Harley, P., Riccio, F., Wu, L., Song, W. and Lieberam, I. (2023) A scalable human iPSC-based neuromuscular disease model on suspended biobased elastomer nanofiber scaffolds. Biofabrication, 15, 045020.

30. Gao, C., Shi, Q., Pan, X., Chen, J., Zhang, Y., Lang, J., Wen, S., Liu, X., Cheng, T.-L. and Lei, K. (2024) Neuromuscular organoids model spinal neuromuscular pathologies in C9orf72 amyotrophic lateral sclerosis. Cell Rep, 43, 113892.

31. Maffioletti, S.M., Sarcar, S., Henderson, A.B.H., Mannhardt, I., Pinton, L., Moyle, L.A., Steele-Stallard, H., Cappellari, O., Wells, K.E., Ferrari, G., et al. (2018) Three-Dimensional Human iPSC-Derived Artificial Skeletal Muscles Model Muscular Dystrophies and Enable Multilineage Tissue Engineering. Cell Reports, 23, 899–908.

32. Lopes, I., Altab, G., Raina, P. and De Magalhães, J.P. (2021) Gene Size Matters: An Analysis of Gene Length in the Human Genome. Front. Genet., 12, 559998.

33. Meyer, L.C. and Wright, N.T. (2013) Structure of giant muscle proteins. Front. Physiol., 4.

34. Savarese, M., Välipakka, S., Johari, M., Hackman, P. and Udd, B. (2020) Is Gene-Size an Issue for the Diagnosis of Skeletal Muscle Disorders? JND, 7, 203–216.

35. Pinheiro, H., Pimentel, M.R., Sequeira, C., Oliveira, L.M., Pezzarossa, A., Roman, W. and Gomes, E.R. (2021) mRNA distribution in skeletal muscle is associated with mRNA size. Journal of Cell Science, 134, jcs256388.

36. Sahakyan, A.B. and Balasubramanian, S. (2016) Long genes and genes with multiple splice variants are enriched in pathways linked to cancer and other multigenic diseases. BMC Genomics, 17, 225.

37. Lim, W.F. and Rinaldi, C. (2023) RNA Transcript Diversity in Neuromuscular Research. J Neuromuscul Dis, 10, 473–482.

38. Logsdon, G.A., Vollger, M.R. and Eichler, E.E. (2020) Long-read human genome sequencing and its applications. Nat Rev Genet, 21, 597–614.

