## Supplementary Information Other for "Identifying fundamental properties of neuromuscular disorder genes through genomic feature analysis"

**Martin *et al.* Genomic features of neuromuscular disorder genes reveal shared and subgroup-specific properties**

#### Supplementary Methods

##### *Gene filtering and harmonization*

Gene entries lacking valid gene symbols and mapping to contigs or pseudo-autosomal regions were removed. Where multiple Ensembl entries were present for the same gene, the entry with the highest transcript count was retained and, in the event of ties, the entry with the highest GC content was selected. All data processing and downstream analyses were performed in R (v.4.3.3).

##### *Gene complexity features*

Features in the gene complexity category, including core gene features, transcripts, exons, homology, and repetitive elements, were defined using genomic annotations from Ensembl v.114 and prior studies, as described below. Transcript-level data included the number of unique transcripts and exons, transcript lengths, and GC content. Canonical transcripts were used to determine the representative transcript length and exon count, and summary isoform statistics (mean, median, minimum, and maximum) were calculated across all genes. We also determined the mean lengths of the 3’ and 5’ UTRs for each gene using Ensembl transcript coordinates. To assess evolutionary conservation, we obtained pairwise percent identity values for orthologs across 11 species. These included non-human primates (*Pan troglodytes*, *Gorilla gorilla*, *Pongo abelii*, *Macaca mulatta*, and *Callithrix jacchus*), as well as well-studied model organisms (*Mus musculus*, *Gallus gallus*, *Xenopus laevis*, *Danio rerio*, *Drosophila melanogaster*, and *Caenorhabditis elegans*). A mean homology score was then calculated across available orthologs for each gene for these species. Paralogy was also assessed by retrieving the number of unique human paralogs associated with each gene from Ensembl. Finally, to characterize repetitive sequence content, we incorporated STR annotations (1), including total STR counts, counts by motif length (i.e., 1-6 bp), and regional counts (i.e., exonic, 5’ UTR, 3’ UTR) for each gene.

##### *Genetic variation features*

To define genetic variation features associated with genes, we incorporated measures of population-level variant tolerance and evolutionary selection. To evaluate the genetic constraint of each gene in human populations, we incorporated *o/e* variant ratios for LoF, missense, and synonymous variants from gnomAD v.2.1.1, which includes data from 125,748 exomes and 15,708 genomes (2). Lower *o/e* ratios indicate greater intolerance to the specific type of variation. Genes with multiple gnomAD entries were filtered to retain those with the lowest LoF *o/e* ratio, followed by the lowest missense *o/e* ratio, ensuring the most constrained gene entry was captured. Genes under positive selection in humans were then identified using data from a study that examined protein-coding genes involved in brain development and cognition across primate evolution (3). Their comparative analysis included data from non-human primates, ancient hominins (e.g., Denisovans), and modern humans to detect lineage-specific evolutionary signals. Gene-level annotations based on omega ratios ($\omega = d_N/d_S$) were used to quantify selection pressure, with $\omega < 1$ indicating negative selection, $\omega > 1$ indicating positive selection, and $\omega = 1$ indicating neutral evolution.

***Gene expression features***

We accessed publicly available transcriptomic datasets to obtain features related to tissue specificity, temporal dynamics, and central nervous system (CNS) relevance, capturing gene expression-related metrics. We incorporated tissue-specific expression data from GTEx v.10 project (4), including *t*-statistics for 53 unique tissues (5). To quantify temporal patterns of gene expression, we also extracted features from a dataset (6), which profiled human organ transcriptomes at 23 time points spanning from four weeks post-conception to 60 years of age using bulk RNA sequencing. From this dataset, we extracted three types of features: (i) metrics indicating whether a gene showed significant dynamic expression over time in a given tissue, based on maSigPro clustering; (ii) indicators of whether a gene was classified as DDG, defined by significant expression changes between fetal and adult stages; and (iii) *Tau* scores, which range from 0 to 1 and quantify tissue specificity, with higher values indicating more restricted expression to a particular tissue. Lastly, to capture cell-type specificity within the CNS, we included fidelity scores from a study (7) that characterized gene expression across major human brain cell types. These scores quantify the extent to which gene expression is restricted to a major CNS cell type, including astrocytes, oligodendrocytes, microglia, and neurons.

***Gene trait features***

Finally, we captured functional information under the gene traits category relevant to understanding genes involved in conditions with a neurological phenotype, such as NMD. These included features relating to neuronal survival and functional gene essentiality in the CNS, derived from both mouse (*Mus musculus*) and human model systems. Mouse CNS essential genes were identified using *in vivo*, genome-wide shRNA and CRISPR knockdown screens (8), which measured the impact of gene knockdown on neuronal survival during brain development. For each gene, we extracted normalized *Z*-scores as indicators of essentiality, where more negative values reflect greater reductions in neuronal viability and suggest greater functional importance. When multiple entries existed for a gene, we retained the entry with the lowest *Z*-score. Mouse gene symbols were then mapped to their human orthologs using Ensembl, via the *biomaRt* package, to enable integration with our human protein-coding gene dataset. We also incorporated neuronal survival data from CRISPR-based functional genomics screens in human iPSC-derived neurons, with and without antioxidant treatment (9). Phenotype scores were included from both CRISPR interference (CRISPRi) and CRISPR activation (CRISPRa) conditions, with the most significant replicate retained per gene (i.e., lowest *P*-value, strongest effect).

#### Supplementary Results

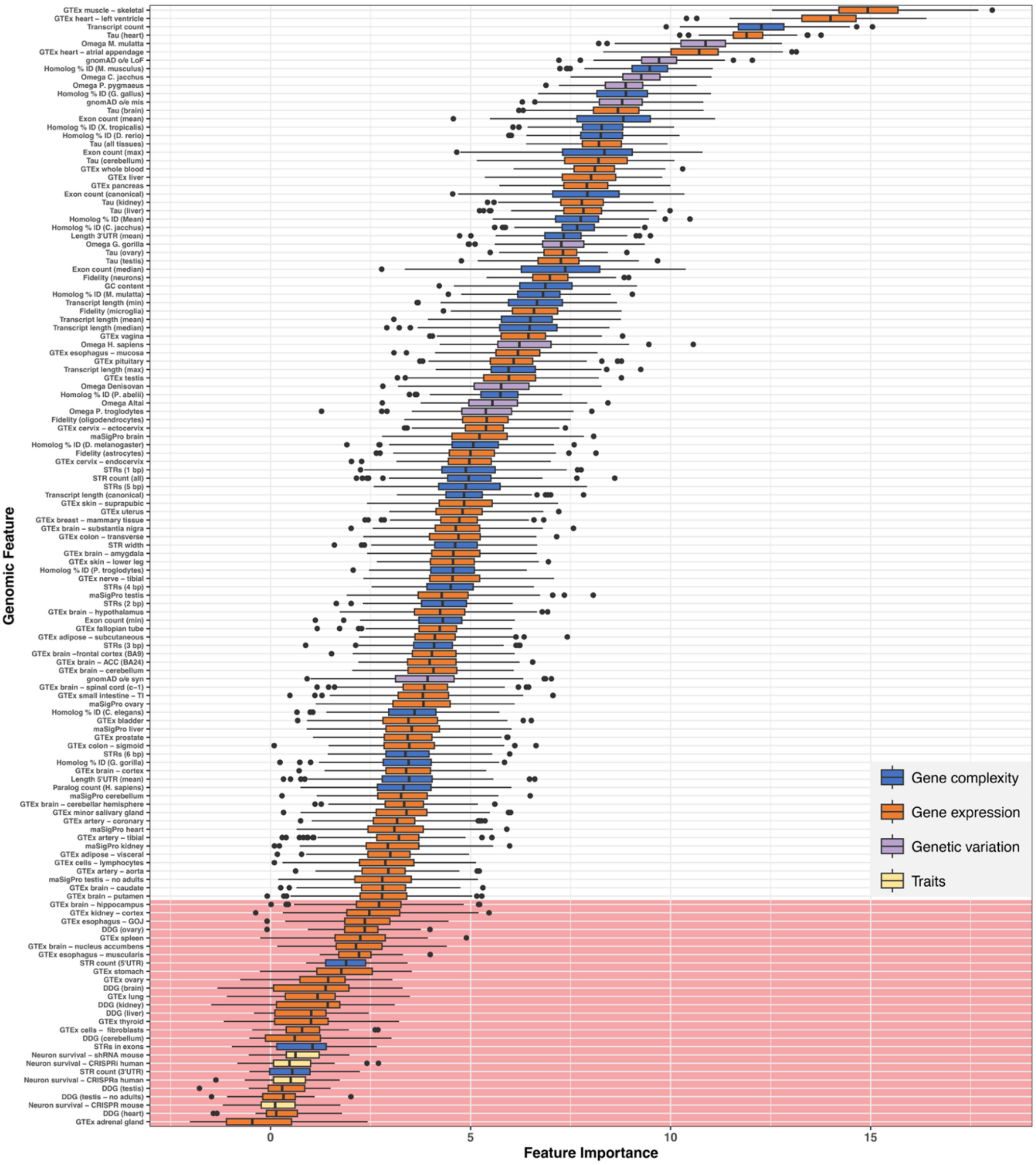


**Supplementary Figure S1. Boruta feature-selection results across all genomic features.** Boruta-based feature selection results ranked by importance scores for 134 curated genomic features, grouped into four categories: gene complexity (*n* = 38, blue), expression (*n* = 81, orange), genetic variation (*n* = 11, lavender), and gene traits (*n* = 4, pale yellow). Features are ranked from most to least important (top to bottom) in differentiating NMD genes (*n* = 639) from all other autosomal protein-coding genes (*n* = 17,880). Features rejected by Boruta are shown at the bottom of the plot and highlighted by red shading. Randomized shadow features were included in the Boruta analysis but are not shown in the figure.

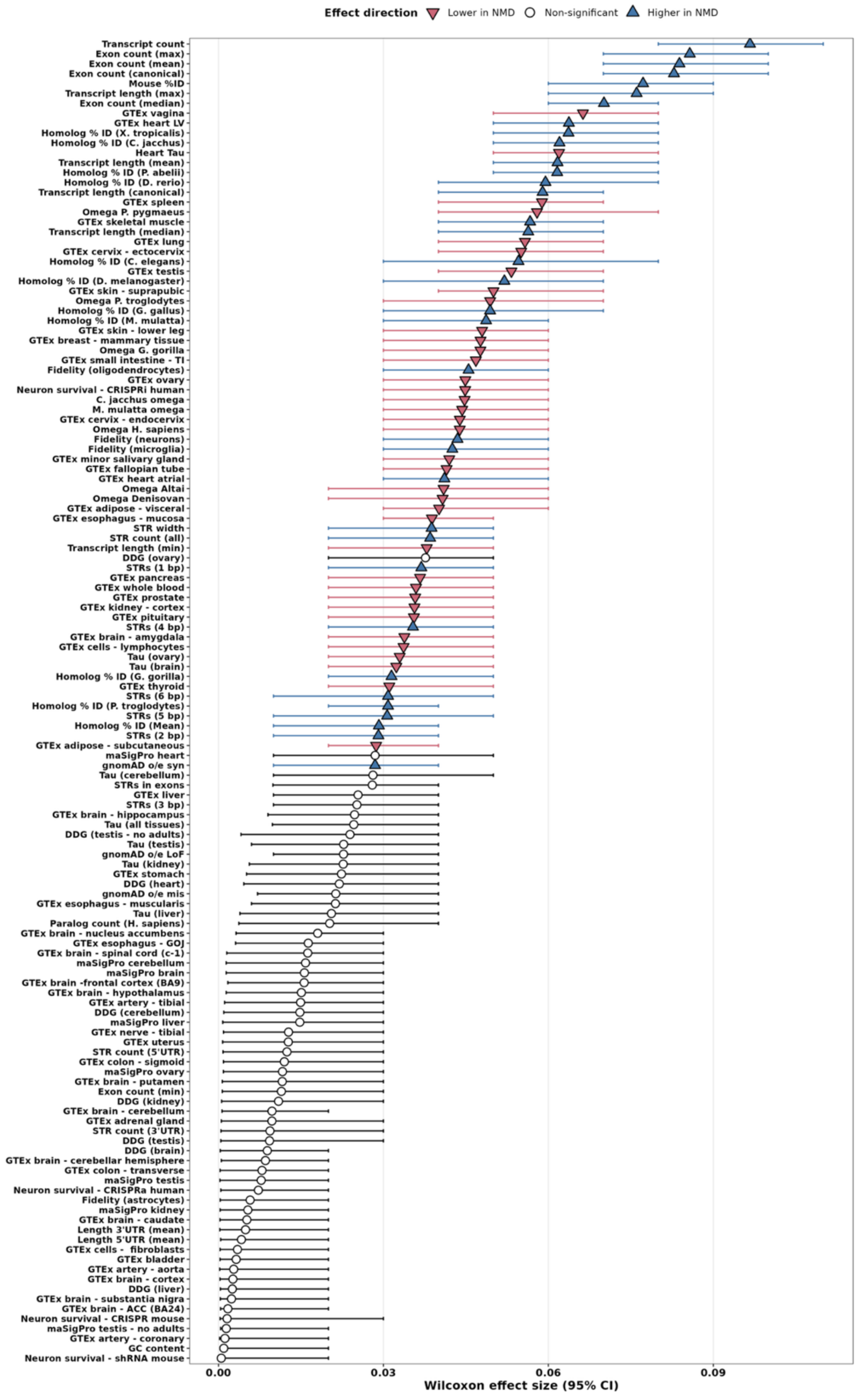


**Supplementary Figure S2. Wilcoxon effect sizes for genomic features distinguishing NMD genes from other autosomal protein-coding genes.** Absolute Wilcoxon effect sizes and 95% confidence intervals are shown for genomic features comparing NMD genes (*n* = 639) with other autosomal protein-coding genes (*n* = 17,880). Features are ordered by effect-size magnitude. Blue upward triangles indicate features with higher values in NMD genes, rose downward triangles indicate lower values in NMD genes, and open circles indicate features that were not significant after Bonferroni correction.

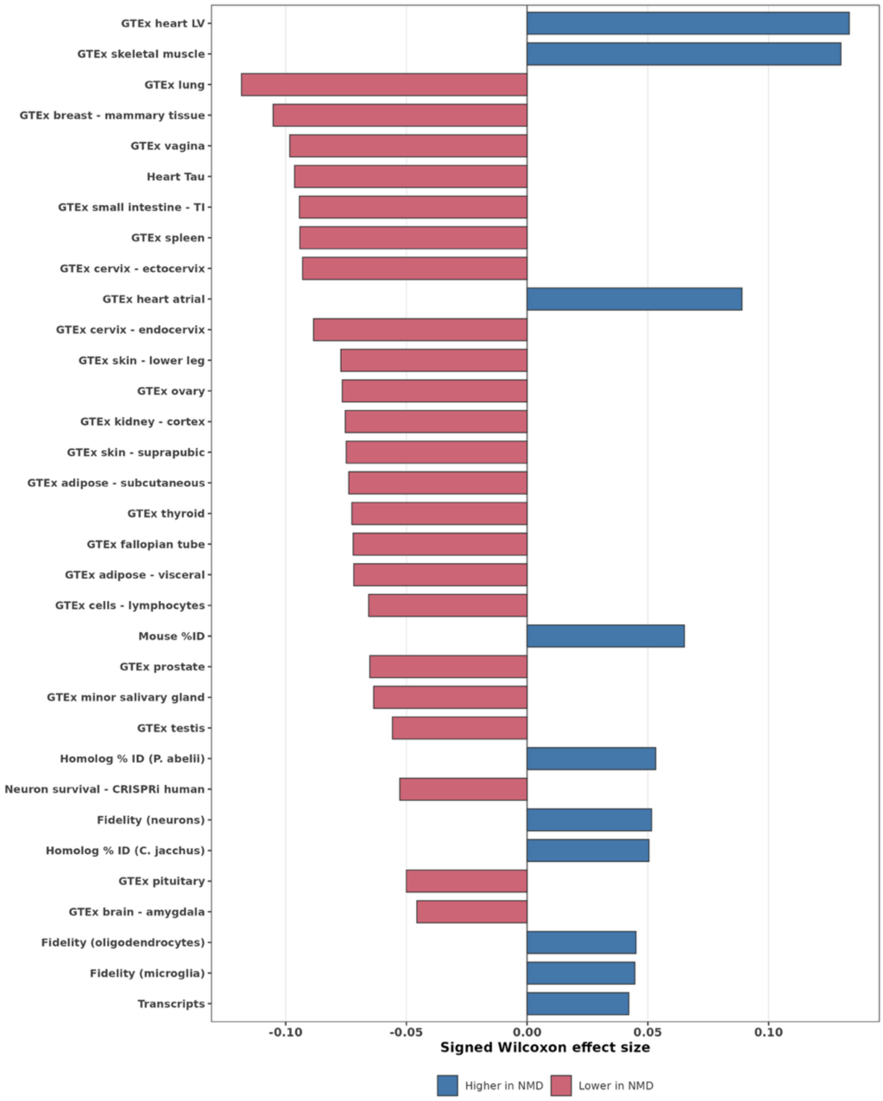


**Supplementary Figure S3. Genomic features that distinguish NMD genes from other high-confidence disease genes.** Signed Wilcoxon effect sizes are shown for the 33 genomic features meeting the FDR-based NMD-distinguishing criteria. Features were required to differ significantly between NMD genes ($n$ = 639) and high-confidence GenCC non-NMD disease genes ($n$ = 3,468), to differ significantly between NMD genes and other autosomal protein-coding genes, and to show a concordant direction of effect across both comparisons (Benjamini-Hochberg FDR < 0.05). Positive Wilcoxon $r$ values (blue) indicate higher feature values in NMD genes and negative values (rose) indicate lower feature values in NMD genes. High-confidence GenCC genes were defined as those with Definitive or Strong gene-disease classifications after excluding NMD genes. Abbreviations: CRISPRi, CRISPR interference; GTEx, Genotype-Tissue Expression; LV, left ventricle; UTR, untranslated region.

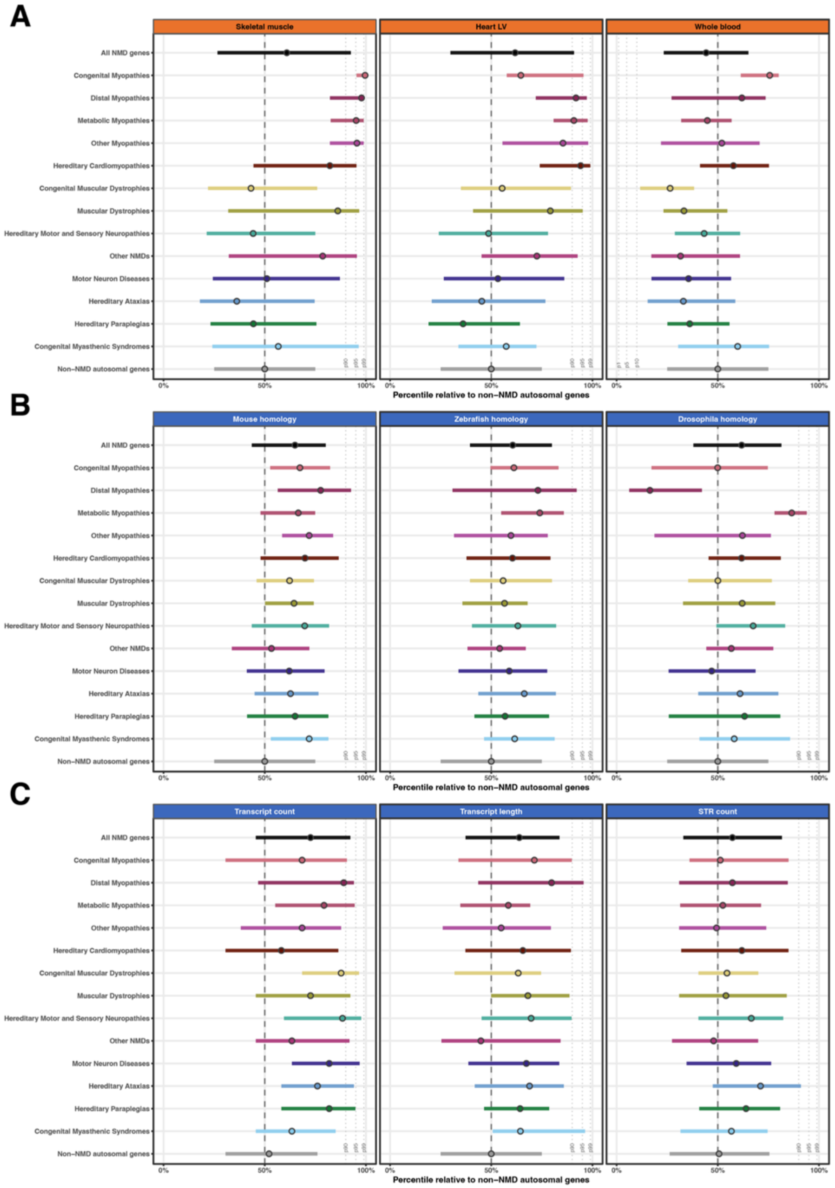

A
Skeletal muscle
Heart LV
Whole blood
All NMD genes
Congenital Myopathies
Distal Myopathies
Metabolic Myopathies
Other Myopathies
Hereditary Cardiomyopathies
Congenital Muscular Dystrophies
Muscular Dystrophies
Hereditary Motor and Sensory Neuropathies
Other NMDs
Motor Neuron Diseases
Hereditary Ataxias
Hereditary Paraplegias
Congenital Myasthenic Syndromes
Non−NMD autosomal genes
0%
50%
100%
Percentile relative to non−NMD autosomal genes
B
Mouse homology
Zebrafish homology
Drosophila homology
C
Transcript count
Transcript length
STR count

**Supplementary Figure S4. Selected expression, homology, and gene complexity features across NMD disease groups.** Median feature percentiles and interquartile ranges are shown for 13 NMD disease groups containing at least 10 genes, all NMD genes, and non-NMD autosomal genes. Feature values were converted to percentiles relative to non-NMD autosomal genes. **(A)** GTEx skeletal muscle, heart left ventricle, and whole-blood expression. **(B)** Homolog percent identity with *Mus musculus* (mouse), *Danio rerio* (zebrafish), and *Drosophila melanogaster* (fruit fly). **(C)** Transcript count, mean transcript length, and total STR count. Dashed vertical lines indicate the 50th percentile. Dotted lines indicate the 90th, 95th, and 99th percentiles for features with higher values in NMD genes or the 1st, 5th, and 10th percentiles for features with lower values in NMD genes. Genes associated with more than one disease group were retained in each relevant group. Abbreviations: GTEx, Genotype-Tissue Expression; LV, left ventricle; STR, short tandem repeat.

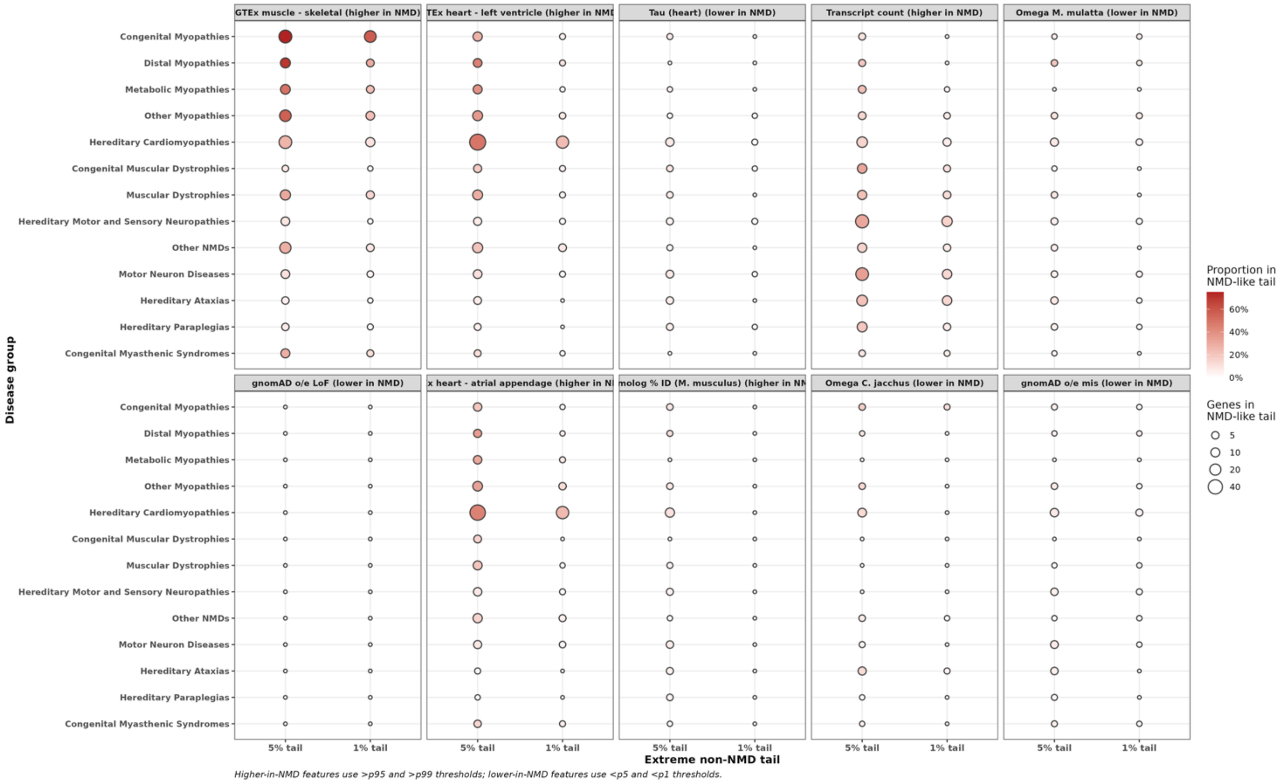


**Supplementary Figure S5. NMD disease groups differ in the proportion of genes with extreme NMD-like feature values.** The proportion and number of genes within direction-specific extreme tails are shown for the 10 highest-ranked Boruta features across NMD disease groups. For features with higher values in NMD genes, the 5% and 1% tails correspond to values above the 95th and 99th percentiles of non-NMD autosomal genes, respectively. For features with lower values in NMD genes, the corresponding tails comprise values below the 5th and 1st percentiles. Point colour indicates the proportion of genes within the NMD-like tail, and point size indicates the number of genes in that tail. Proportions were calculated using genes with non-missing values for the corresponding feature as the denominator. Genes associated with more than one disease group were retained in each relevant group. Abbreviations: GTEx, Genotype-Tissue Expression; LV, left ventricle; LoF, loss-of-function; mis, missense; o/e, observed/expected.

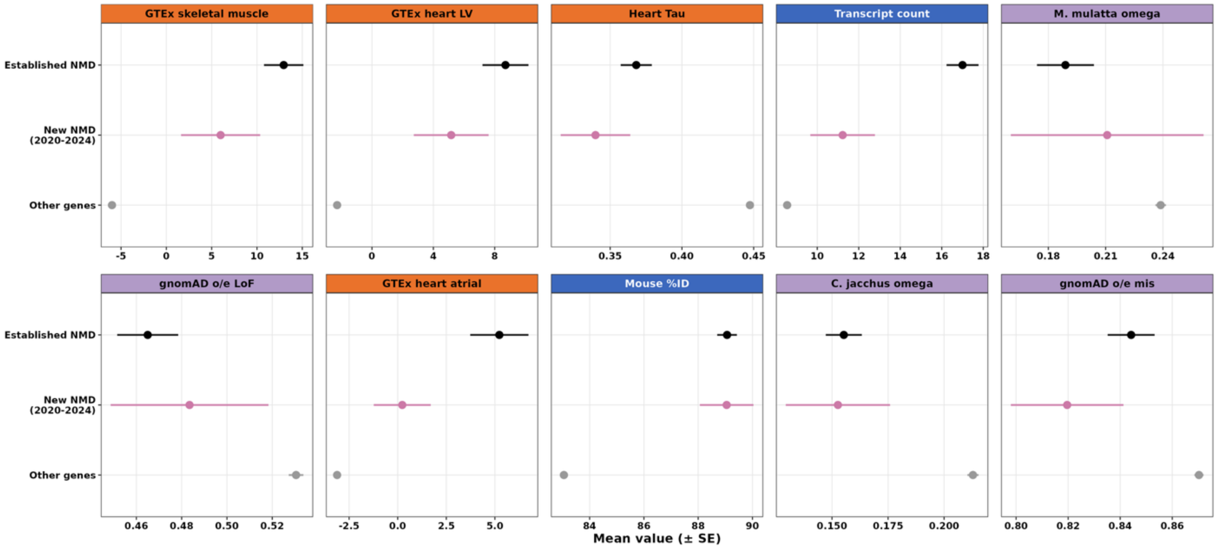


**Supplementary Figure S6. Top-ranked genomic features in established and more recently added NMD genes.** Mean values (± standard error, SE) are shown for the 10 highest-ranked Boruta features in NMD genes established by July 2020 ($n$ = 550), genes added to the NMD gene list between 2020 and 2024 ($n$ = 89), and other autosomal protein-coding genes ($n$ = 17,880). Group sizes indicate the number of genes assigned to each group; plotted means and SEs use available values for each feature. Established NMD genes are shown in black, more recently added NMD genes in rose, and other autosomal protein-coding genes in grey. Abbreviations: GTEx, Genotype-Tissue Expression; LV, left ventricle; LoF, loss-of-function; mis, missense; o/e, observed/expected; SE, standard error.

**Supplementary Table 2.** NMD subcategories as described in the GeneTable of Neuromuscular Disorders (www.musclegenetable.fr; Benarroch *et al.*, 2024).

| Disease Group | Example(s) | Number of Genes Identified |
|---|---|---|
| 1. Muscular dystrophies | Duchenne/Becker muscular dystrophies, Emery-Dreifuss muscular dystrophies, limb girdle muscular dystrophies | 53 |
| 2. Congenital muscular dystrophies | Congenital muscular dystrophies, Bethlem myopathies, muscular dystrophy-dystroglycanopathies | 38 |
| 3. Congenital myopathies | Congenital myopathies, nemaline myopathies, centronuclear myopathies | 41 |
| 4. Distal myopathies | Miyoshi muscular dystrophies, distal myopathies | 22 |
| 5. Other myopathies | Myofibrillar myopathies, oculopharyngodistal myopathies | 45 |
| 6. Myotonic syndromes | Myotonic dystrophies, myotonias, rippling muscle diseases | 6* |
| 7. Ion channel muscle diseases | Chloride/sodium/potassium/calcium channel muscle diseases | 8* |
| 8. Malignant hyperthermia | Malignant hyperthermia susceptibilities | 2* |
| 9. Metabolic myopathies | Glycogen storage diseases, glycolytic pathway diseases, disorders of lipid metabolism | 29 |
| 10. Hereditary cardiomyopathies | Hypertrophic/dilated/restrictive/arrhythmogenic ventricular/other non-arrhythmogenic hereditary cardiomyopathies, long/short QT syndromes, atrial fibrillation syndromes | 116 |
| 11. Congenital myasthenic syndromes | Slow-channel/fast-channel/acetylcholine receptor congenital myasthenic syndromes | 35 |

| | | | |
|---|---|---|---|
| 12. | Spinal muscular atrophies motoneuron diseases | Proximal spinal muscular atrophies, distal hereditary motor neuropathies, amyotrophic lateral sclerosis | 94 |
| 13. | Hereditary ataxias | Spinocerebellar ataxias, episodic ataxias | 90 |
| 14. | Hereditary motor and sensory neuropathies | Charcot-Marie-Tooth neuropathies, Dejerine-Sottas syndromes, other hereditary sensory and autonomic neuropathies, other complex neuropathy syndromes | 115 |
| 15. | Hereditary paraplegias | Spastic paraplegias, spastic ataxias | 76 |
| 16. | Other neuromuscular disorders | Torsion/myotonic dystonias, fibrosis of extraocular muscles, distal arthrogryposis, fetal akinesia deformation sequences, progressive external opthalmoplegias, mitochondrial DNA depletion syndromes, etc. | 75 |

*Not included in sub-group analyses due to <10 genes.

### References


1. Chen, Z., Tucci, A., Cipriani, V., Gustavsson, E.K., Ibañez, K., Reynolds, R.H., Zhang, D., Vestito, L., García, A.C., Sethi, S., *et al.* (2023) Functional genomics provide key insights to improve the diagnostic yield of hereditary ataxia. *Brain*, **146**, 2869–2884.
2. Chen, S., Francioli, L.C., Goodrich, J.K., Collins, R.L., Kanai, M., Wang, Q., Alföldi, J., Watts, N.A., Vittal, C., Gauthier, L.D., *et al.* (2024) A genomic mutational constraint map using variation in 76,156 human genomes. *Nature*, **625**, 92–100.
3. Dumas, G., Malesys, S. and Bourgeron, T. (2021) Systematic detection of brain protein-coding genes under positive selection during primate evolution and their roles in cognition. *Genome Res*, **31**, 484–496.
4. GTEx Consortium (2017) Genetic effects on gene expression across human tissues. *Nature*, **550**, 204–213.
5. Finucane, H.K., Reshef, Y.A., Anttila, V., Slowikowski, K., Gusev, A., Byrnes, A., Gazal, S., Loh, P.-R., Lareau, C., Shoresh, N., *et al.* (2018) Heritability enrichment of specifically expressed genes identifies disease-relevant tissues and cell types. *Nat Genet*, **50**, 621–629.
6. Cardoso-Moreira, M., Halbert, J., Valloton, D., Velten, B., Chen, C., Shao, Y., Liechti, A., Ascenção, K., Rummel, C., Ovchinnikova, S., *et al.* (2019) Gene expression across mammalian organ development. *Nature*, **571**, 505–509.
7. Kelley, K.W., Nakao-Inoue, H., Molofsky, A.V. and Oldham, M.C. (2018) Variation among intact tissue samples reveals the core transcriptional features of human CNS cell classes. *Nat Neurosci*, **21**, 1171–1184.
8. Wertz, M.H., Mitchem, M.R., Pineda, S.S., Hachigian, L.J., Lee, H., Lau, V., Powers, A., Kulicke, R., Madan, G.K., Colic, M., *et al.* (2020) Genome-wide In Vivo CNS Screening Identifies Genes that Modify CNS Neuronal Survival and mHTT Toxicity. *Neuron*, **106**, 76-89.e8.
9. Tian, R., Abarientos, A., Hong, J., Hashemi, S.H., Yan, R., Dräger, N., Leng, K., Nalls, M.A., Singleton, A.B., Xu, K., *et al.* (2021) Genome-wide CRISPRi/a screens in human neurons link lysosomal failure to ferroptosis. *Nat Neurosci*, **24**, 1020–1034.
